# SERINC5 Is the Primary Target of Retroviral Antagonists Among Human SERINC Paralogs and Requires a Tri-Cysteine Motif for Its Counteraction by HIV-1 Nef

**DOI:** 10.1101/2025.10.27.684794

**Authors:** Cinzia Bertelli, Annachiara Rosa, Teresa Vanzo, Ajit Chande, Remigiusz Serwa, Edward Tate, Peter Cherepanov, Massimo Pizzato

**Affiliations:** Department of Cellular, Computational and Integrative Biology, University of Trento, Italy; Chromatin Structure and Mobile DNA Laboratory, Francis Crick Institute, London, UK; Department of Chemistry, Imperial College London, London, UK; Department of Infectious Disease, Imperial College London, London, UK

## Abstract

The human transmembrane protein SERINC5 incorporates into budding HIV-1 particles and inhibits their fusion with new target cells. Among the five members of the SERINC family, SERINC3 and SERINC1 exhibit modest antiviral activity, whereas SERINC2 lacks this function. HIV-1 counteracts SERINC5 by promoting its endocytosis in a Nef-dependent manner. SERINC5 is also downregulated by glycoGag of Moloney murine leukaemia virus (MoMLV) and by the S2 glycoprotein of equine infectious anaemia virus (EIAV).

Here, we demonstrate that all human SERINCs, despite their differing antiviral potencies, are incorporated into HIV-1 virions. SERINC5 is the most sensitive to viral antagonists, whereas its paralogs display moderate (SERINC2, SERINC3) or no (SERINC1) susceptibility, suggesting that SERINC5 has exerted stronger selective pressure on retroviral evolution.

To identify the determinants of SERINC5 sensitivity to Nef, we performed structure-guided mutagenesis and identified a critical tri-cysteine motif (Cys355, Cys356, Cys358) within intracellular loop 4 (ICL4) required for Nef-mediated downregulation. Substitution of these residues with glutamines conferred resistance to Nef antagonism. While this motif is essential for SERINC5 internalization by diverse HIV-1 and SIV Nef alleles, it is dispensable for downregulation by MoMLV glycoGag and EIAV S2, indicating that these viral antagonists exploit distinct structural features of SERINC5. Finally, we show that SERINC5 is palmitoylated at the tri-cysteine motif, raising the possibility that this modification modulates its sensitivity to Nef.

Together, these results confirm the central role of SERINC5 as a target of retroviral counteraction and highlight the crucial importance of ICL4 in mediating its engagement by Nef.

**Importance:** The SERINC family of multipass transmembrane proteins includes host factors capable of restricting retrovirus infectivity. Among the five human paralogs, SERINC5 displays the strongest antiviral activity and is the only member efficiently targeted by all known retroviral antagonists, including HIV-1 Nef, MLV glycoGag, and the EIAV S2. This selectivity indicates that the evolutionary pressure exerted by SERINC proteins on retroviruses has been primarily driven by SERINC5 rather than by its relatives. Through structure-guided mutagenesis, we identified a tri-cysteine motif within intracellular loop 4 as a crucial determinant for SERINC5 downregulation by Nef, providing a molecular explanation for its distinctive susceptibility. These findings establish SERINC5 as the principal target of viral counteraction and suggest that its molecular features have contributed to shaping the adaptation of primate lentiviral Nef proteins to the host.

## Introduction

The SERINC family of multipass transmembrane proteins is highly conserved across eukaryotic species, with orthologs identified in organisms ranging from unicellular eukaryotes to mammals^1^. The family includes five human paralogs (SERINC1-5), among which SERINC5 was identified as a potent inhibitor of HIV-1 infectivity^2,3^. Human SERINC5 incorporates into budding HIV-1 virions and restricts the early stages of viral life cycle by interfering with the virus-cell fusion process^4–6^. Remarkably, this antiviral activity is not unique to human SERINC5, as it is conserved across SERINC orthologs from distantly related species such as *Drosophila melanogaster*^7–11^, suggesting that the ability to inhibit retroviral infection relies on a fundamentally conserved feature of this protein family. Consistent with this evolutionary conservation, human SERINC paralogs share a characteristic molecular architecture^8^, though they inhibit retrovirus infection with different potencies^2,7,12^. While SERINC5 demonstrates the most robust restriction of retroviral infectivity, SERINC1 and SERINC3 exert weaker antiviral effects, and SERINC2 lacks detectable antiviral function^12,13^. While SERINCs were reported to disrupt the viral membrane asymmetry by acting as lipid scramblases^14^, this activity may not be responsible for the inhibition of retroviral infectivity^15^.

Retroviruses have evolved countermeasures to overcome the potent restriction imposed by SERINC5. In primate lentiviruses, the accessory protein Nef antagonizes SERINC5 by promoting its removal from the plasma membrane through clathrin- and AP2-mediated endocytosis, redirecting it to late endosomes and preventing its incorporation into virions^2,3,16^. Similarly, in Equine Infectious Anaemia Virus (EIAV) and gammaretroviruses, the accessory proteins S2 and glycoGag, respectively, fulfil a comparable function^17,18^.

These independent evolutionary adaptations, arising in three distinct retroviral lineages, suggests that SERINC5 has exerted significant selective pressure on retrovirus evolution. However, the molecular basis of SERINC5 targeting by viral antagonists remains largely unknown. Notably, SERINC5 was recently reported to interfere with viral species outside of the retroviral family^19–22^, potentially underscoring a broader antiviral activity that has driven the emergence of viral antagonists beyond retroviruses.

While SERINC5 has the most potent antiviral activity, other SERINC paralogs may also contribute to the restriction of retroviral infection. SERINC1, SERINC3, and SERINC4 have been reported to exert some degree of antiviral activity, though their potencies are generally weaker than that of SERINC5^2,7,12^. The relevance of the antiviral function of each paralog remains, therefore, a compelling question. Similarly, whether the known viral antagonists have evolved to target all SERINC isoforms, as well as the molecular basis of their antagonistic activity, remain largely unexplored.

Here, we investigated the selectivity with which the known retroviral counteracting factors antagonize the SERINC proteins and observed that Nef, glycoGag and S2 are significantly more active against SERINC5 than against the other paralogs. Furthermore, we identified a crucial molecular determinant in SERINC5 required for the selective counteraction by HIV-1 Nef, indicating a different molecular basis for antagonization by the three retroviral factors.

## Results

### Human SERINC paralogs are differently targeted by HIV-1 Nef

We first investigated the extent of the SERINC paralogs activity against HIV-1 and their counteraction by HIV-1 Nef. Nef from the subtype C isolate 97ZA012 isolate was chosen because it is representative of the primary HIV-1 strains most prevalent in the world and, in previous experiments, has proven to be significantly more active than lab-adapted strains in promoting HIV-1 infectivity in the presence of SERINC5^2^. HIV-1 NL4-3 limited to a single cycle of replication and expressing Nef_97ZA012_ or an inactive mutant (*nef_fs_*) was produced by transfection of SERINC3/5 double knockout HEK293T cells (S3/S5 DKO) with a provirus construct and vectors encoding Env_HXB2_ and SERINC variants. To avoid saturation of Nef antagonism, we expressed *SERINC* paralogs from vectors that contain promoters tuned for moderate (pBJ5) and low (pBJ6) expression in HEK293T cells^2^. The infectivity of the progeny viruses, normalized on their reverse transcriptase (RT) activity, was measured on TZM-zsGreen reporter cells.

Consistent with previous data, SERINC5 caused the most robust inhibition of HIV-1 infectivity (more than 25-fold) even at the lowest expression level obtained with the pBJ6 vector, while SERINC2 had no detectable effect on Nef*-*deficient virions (Figure 1A). Nef_97ZA012_ fully restored HIV-1 infectivity when SERINC5 was expressed from pBJ6, while partially rescuing infectivity in the presence of the higher SERINC5 levels driven by pBJ5 vector. Nef was not required for optimal virus infectivity in the presence of SERINC2, since this paralog did not exert any restrictive activity, in line with previous data^2,7,12^. SERINC1 and SERINC3 had only a modest inhibitory activity (∼ 3-fold) on the *nef*- defective virus, even at their highest expression level. Nevertheless, the inhibitory effect of SERINC3 was completely antagonized by Nef, confirming previous evidence that this paralog is sensitive to the HIV-1 accessory protein^2,3^. By contrast, Nef did not rescue the moderate infectivity defect of HIV-1 caused by the expression of SERINC1, indicating that this paralog may not be susceptible to viral counteraction. Compared to SERINC5, the effect of SERINC4 on HIV-1 infectivity appears lower in magnitude. Nonetheless, Nef increased virus infectivity in the presence of SERINC4, suggesting that Nef targets this paralog.

**Figure 1:**
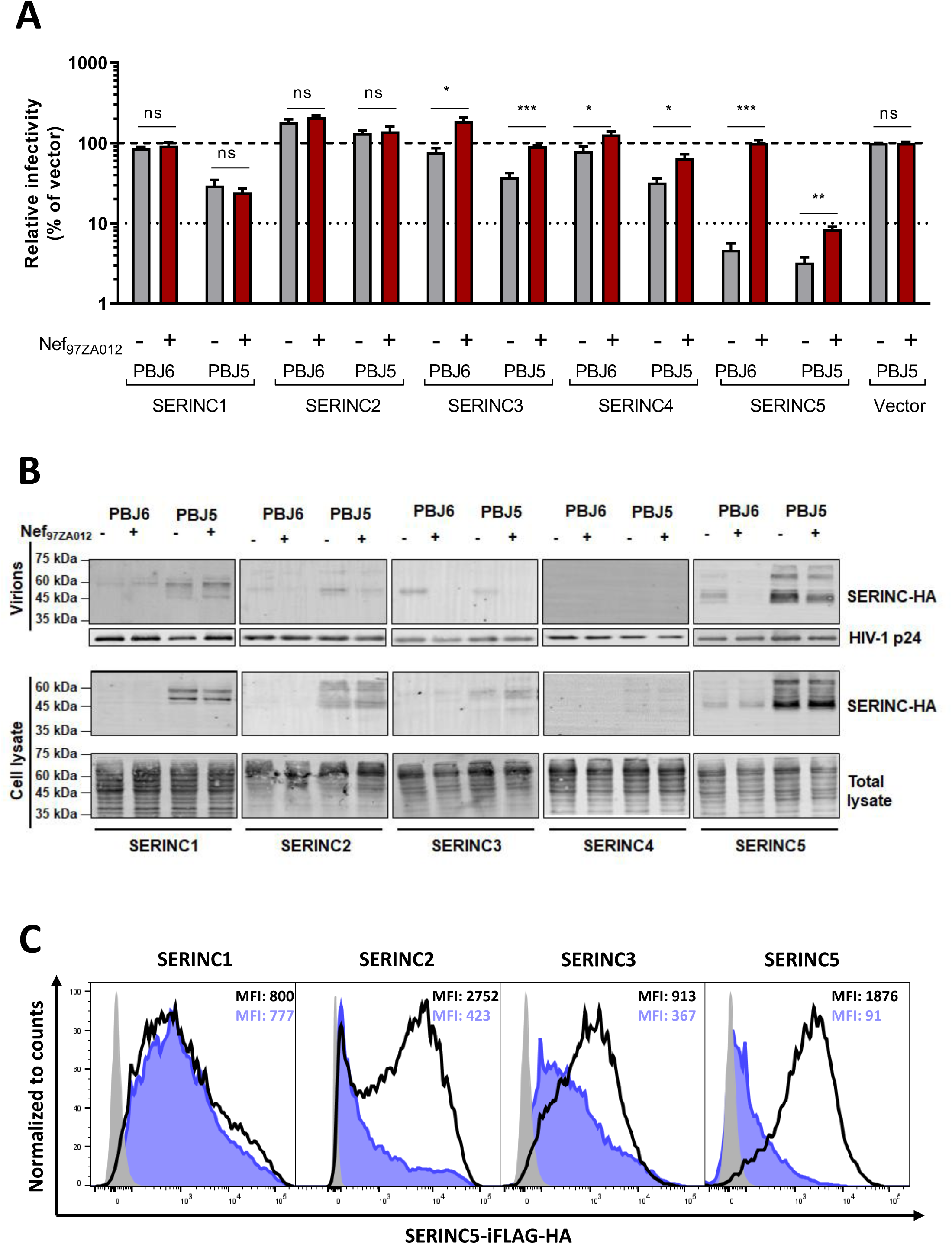
HIV-1 Nef internalizes each human SERINC paralog with diverse efficiency. (A-B) Δenv HIV-1_NL4-3_, encoding either nef_97ZA012_ or nef_fs_, was produced in S3/S5 DKO HEK293T cells, co-transfected with PBJ5-Env_HXB2_ and a human SERINC-HA paralog, expressed from PBJ5 or PBJ6 vector as indicated. (A) The infectivity of progeny virions was measured on TZM-ZsGreen cells and normalized on the RT activity of the viral inoculum. Infectivity is expressed relative to the infectivity of virions produced in the absence of SERINCs. A representative experiment is shown of n=2 biological independent replicates. Bar graphs represent the mean + SD of n=4 technical replicates. Statistical significance was measured with unpaired two-tailed t-test. *p<0.05; **p<0.01; ***p<0.001; ns: not significant; (B) Detection of human SERINCs in purified virus pellets and virus-producer cell lysates by western blot; (C) The efficiency of HIV-1 Nef to promote SERINCs internalization was measured by detecting the amount of SERINCs exposed on the cell surface by flow cytometry. S3/S5 DKO HEK293T cells were co-transfected with pIRES2-eGFP expressing Nef_97ZA012_ and a human SERINC-iFLAG-HA paralog, expressed from pcDNA3.1 plasmid. Cells were labelled with an anti-FLAG antibody and an APC-conjugated secondary antibody. The Median Fluorescence Intensity (MFI) of APC in the transfected GFP+ population is indicated.

Current evidence supports the idea that Nef enhances HIV-1 infectivity by preventing the incorporation of SERINC5 into progeny virions^2,3^. Therefore, we investigated to what extent Nef can prevent the association of each SERINC paralog with virion particles. Viruses and producer cells were analysed by western blotting to detect SERINC proteins in pelleted virus particles and cell lysates (Figure 1B). While SERINC5 and SERINC3 were incorporated into Nef-negative virions, they readily disappeared in virus particles in the presence of Nef, confirming the ability of Nef to counteract the viral antagonists. By contrast, SERINC1 was incorporated into virions irrespective of Nef expression in producer cells, mirroring the inability of Nef to rescue the weak inhibition of infectivity of HIV-1 carrying this SERINC paralog. Despite lacking restriction activity, SERINC2 was incorporated into virions, and the amount of virion-associated protein was decreased in the presence of Nef. This indicates that, despite not interfering with HIV-1 infectivity, SERINC2 can also be targeted by Nef. The expression of SERINC4 resulted in insufficient detection in virions, in line with published evidence that the protein is subject to quick turnover and proteasome-mediates degradation^23^. Given the difficulties in achieving expression levels that can be confidently detected, SERINC4 was excluded from the rest of our investigation.

We and others have shown that Nef prevents the incorporation of SERINC5 into virions by downregulating its expression at the cell surface^2,3,16^. Accordingly, we investigated whether Nef’s ability to exclude SERINC paralogs from virions correlates with its capacity to downregulate their surface expression. Based on a previously established approach^8^, a FLAG epitope was inserted into each SERINC paralog within the extracellular loop 4 (ECL4) to facilitate detection. HIV-1 Nef_97ZA012_ was cloned into a vector upstream of an IRES-eGFP cassette to allow specific gating of Nef-expressing cells in flow cytometry analysis. SERINC-expressing plasmids were co-transfected in S3/S5 DKO HEK293T cells together with a retroviral factor, and the surface expression of SERINC proteins was measured in eGFP-positive cells. While the expression of HIV-1 Nef resulted in a 30-fold downregulation of SERINC5, it reduced SERINC2 surface expression by only 5-fold and SERINC3 expression by approximately 3-fold (Figure 1C). By contrast, SERINC1 expression was not affected by Nef, mirroring the inability of the viral protein to exclude this paralog from virus particles and to counteract the inhibition of infectivity. Collectively, these data indicate that HIV-1 Nef from HIV-1_97ZA012_ can target all human SERINC paralogs, except SERINC1, irrespective of the magnitude of their antiretroviral activity.

### Human SERINC5 is the only paralog readily targeted by all retroviral antagonists

Having established the breadth of the counteracting activity of HIV-1_97ZA012_ Nef towards the different human SERINC paralogs, we investigated the ability of additional *nef* alleles and other retroviral factors to target each human paralog. In addition to Nef from HIV-1_97ZA012_, we selected two other *nef* alleles (derived from HIV-1_SF2_, and SIV_Mac239_), MoMLV glycoGag and EIAV S2. Each of the selected viral factors was expressed upstream of an IRES-eGFP cassette to facilitate gating of transfected cells. To verify the specificity of the counteracting activity, mCAT-1, a multi-span transmembrane protein, was used as a control after inserting an internal FLAG tag within the third extracellular loop. Among the five paralogs, SERINC5 was the only paralog robustly downregulated by all retroviral factors which reduced its expression by 20- to 30-fold (Figures 2A and 2B). SERINC1 was found to be largely resistant to all antagonists, while SERINC2 and SERINC3 were differently targeted by the viral antagonists. Nef variants displayed diverse effects on surface expression of SERINC3 and SERINC2. While Nef_97ZA012_ induced a 2-fold decrease in SERINC3 surface expression (50% residual surface expression), Nef_SF2_ and Nef_Mac2*39*_ did not alter its surface level. Similarly, although SERINC2 was downregulated 5-fold by Nef_97ZA012,_ it was not affected by Nef_SF2_ and Nef_mac239_. MoMLV glycoGag potently downregulated SERINC5, but had a weaker activity against SERINC2 and SERINC3, causing only a 50% decrease in their surface expression. By contrast, EIAV S2 showed the broadest antagonizing activity, causing marked SERINC2 and SERINC3 surface downregulation to a level similar to SERINC5. EIAV S2 also displayed some activity toward SERINC1, though minimal (∼40% residual surface expression) and likely nonspecific, since a comparable effect was observed on the control protein (mCAT-1).

**Figure 2:**
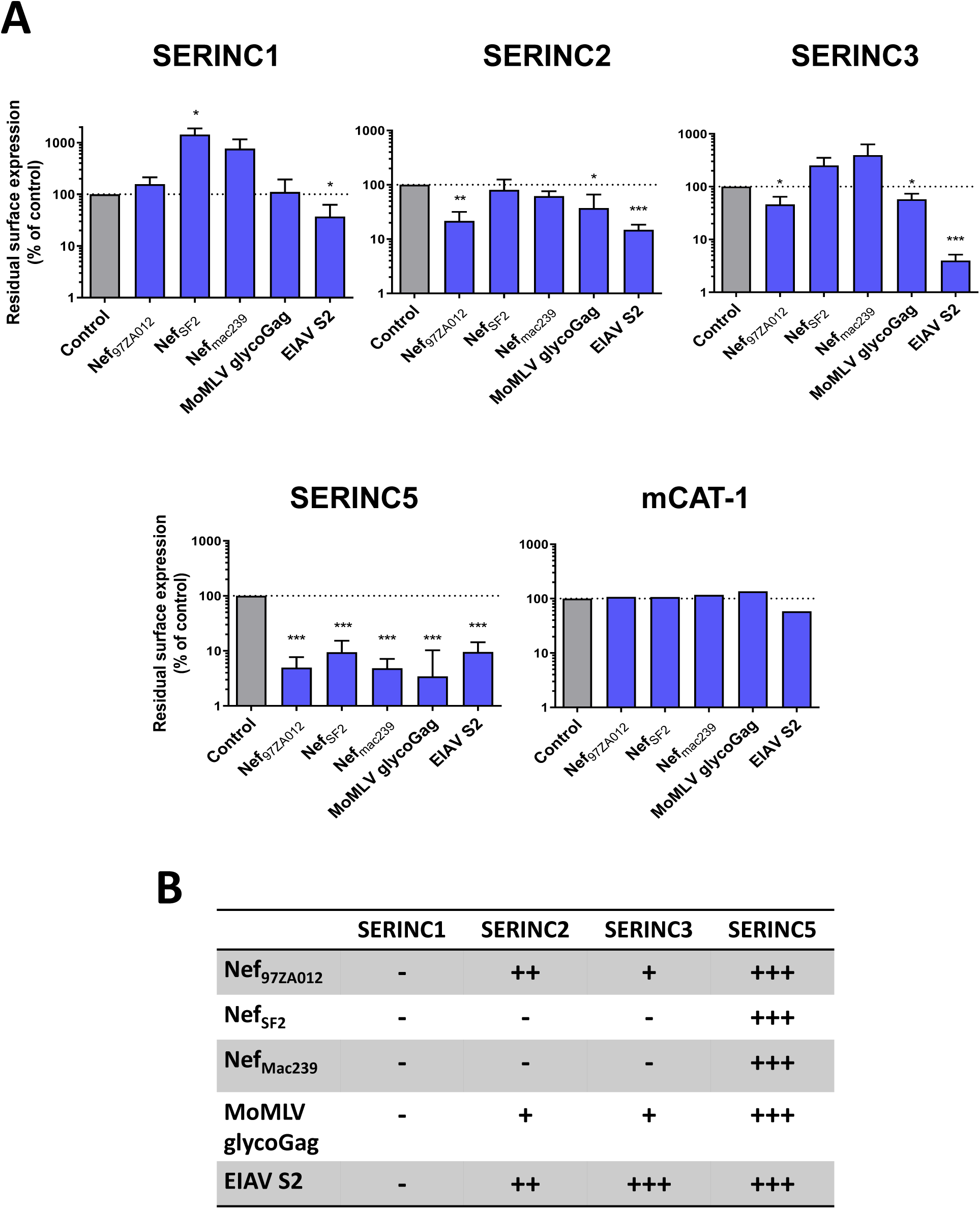
Human SERINC paralogs are differently targeted to internalization by retroviral antagonists. (A) Residual surface expression of each SERINC paralog measured by flow cytometry in the presence of the indicated retroviral antagonists. S3/S5 DKO HEK293T cells were co-transfected with a retroviral factor, expressed from pIRES2-eGFP and a SERINC-iFLAG- HA paralog or, as control, mCAT-1-iFLAG expressed from pcDNA3.1. The residual surface expression in the presence of each retroviral factor was measured as median fluorescence intensity and values are expressed as percentage of the empty control sample, considered as 100% (grey bar). Bar graphs indicate the mean + SD of n=3 biological independent experiments. Significant differences with control (100% residual surface expression) are indicated as follows: *p<0.05; **p<0.01; ***p<0.001 (one-sample t-test); (B) Efficiency of human SERINC paralogs internalization by selected retroviral antagonists, calculated on the residual SERINC surface expression measured in the presence of each viral factor, as follows: ≥50% -; <50% +/-; <30% +; <10% ++.

In conclusion, SERINC5, which exhibits the strongest antiviral activity, is also the paralog most consistently and effectively targeted by all viral antagonists. Conversely, the activity of these viral factors against other SERINC isoforms is more variable and generally weaker. These findings underscore the capacity of viral antagonists such as Nef to discriminate among SERINC family members. Furthermore, the differential downregulation of SERINC paralogs by Nef, glycoGag, and S2 suggests that these factors recognize distinct molecular features within the SERINC proteins to mediate their antagonism.

### Residues within ICL4 are crucial for the sensitivity of SERINC5 to internalisation by HIV-1 Nef

The features of SERINC5 governing its susceptibility to Nef, glycoGag and S2 remain poorly understood. Previous studies have pointed to a crucial role of the fourth intracellular loop, which contains two hydrophobic amino acids (Leu350 and Ile352) required for efficient downregulation by HIV-1 Nef ^10^ and an acidic motif (^364^EDTEE^368^)^24^, found to alter SERINC5 sensitivity to counteraction by Nef. To identify residues important for the susceptibility of the host factor to downregulation by HIV-1 Nef_97ZA012_, we took advantage of the extensive collection of SERINC5 variants that we had previously generated^8^. All SERINC5 mutants within the panel have been previously characterized^8^, allowing us to focus on those that did not interfere with cell surface expression. In addition, our investigation included the SERINC5 variant LI350,352AA, previously studied by Göttlinger and colleagues^10^. As shown in Figure 3A, which displays the residual cell surface protein levels measured upon Nef expression, none of the variants analysed was individually sufficient to completely abrogate SERINC5 susceptibility to downregulation by Nef_97ZA012_. While the viral factor reduced wild-type SERINC5 surface expression to 10%, most substitutions did not substantially impair downregulation by Nef_97ZA012_, as residual surface levels remained between 5 and 20%.

**Figure 3:**
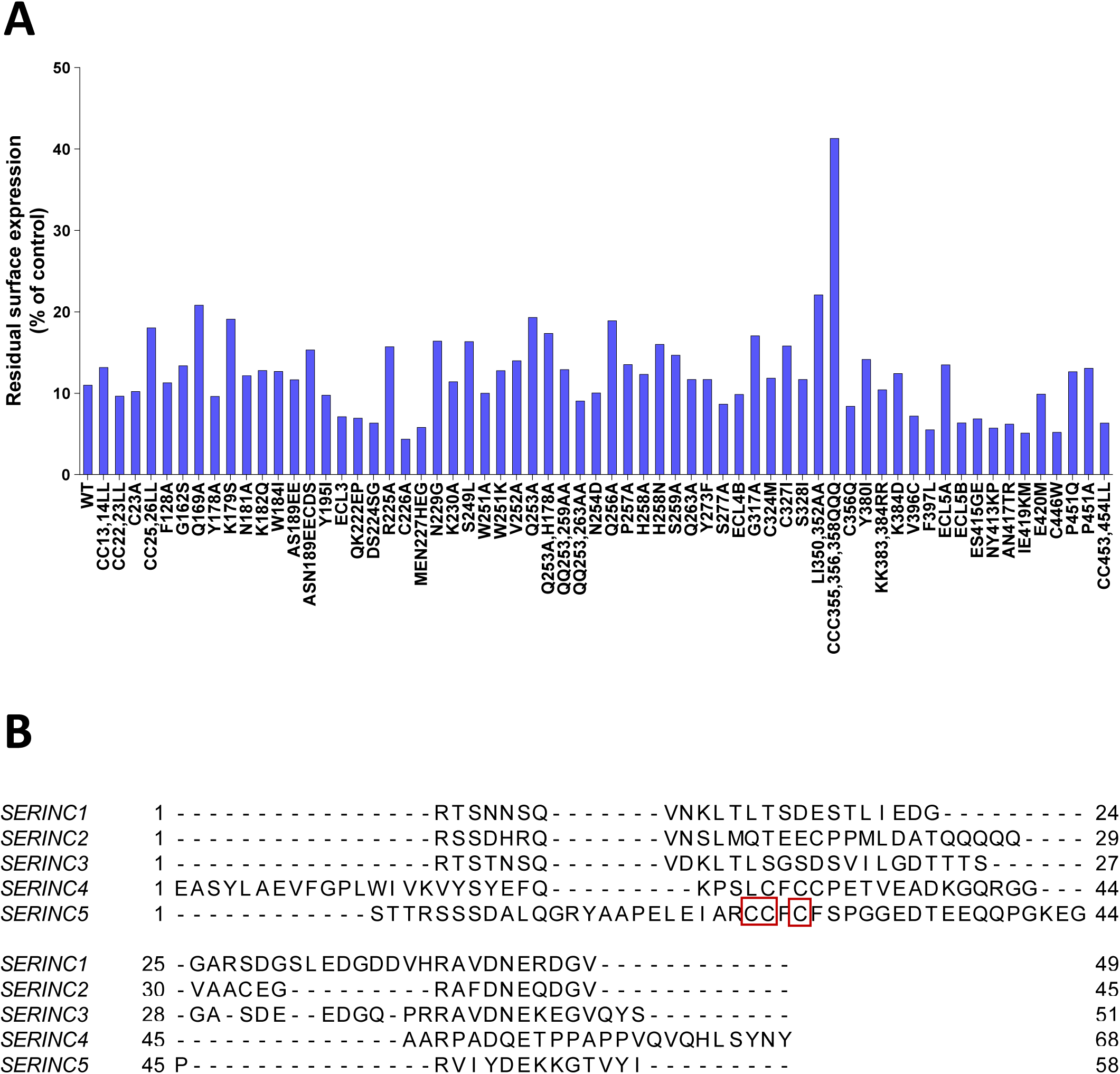
A cluster of cysteine within ICL4 is a major determinant of SERINC5 sensitivity to internalization by HIV-1 Nef_97ZA012_. (A) Sensitivity of a panel of SERINC5 variants to downregulation by Nef_97ZA012_, measured by flow cytometry and calculated as residual surface expression in the presence of the viral factor relative to the empty control sample, considered as 100%. S3/S5 DKO HEK293T cells were co-transfected with a SERINC5-iFLAG-HA variant, expressed from pcDNA3.1, and pIRES2-eGFP-Nef_97ZA012_ or empty vector control. The residual surface expression in the presence of Nef_97ZA012_ was measured as median fluorescence intensity, and values are expressed as a percentage of the empty control sample, considered as 100%; (B) Alignment of ICL4 amino acid sequences of human SERINC paralogs. SERINC proteins topology was analysed with TMHMM2.0 to identify transmembrane regions; the amino acid sequences corresponding to ICL4 were selected and aligned by Clustalω, using default parameters. SERINC5 cysteine stretch within ICL4 is highlighted in red.

However, one variant appeared to stand out by acquiring greater resistance to Nef_97ZA012_- mediated internalization (more than 40% residual expression). In this variant, three contiguous cysteine residues (Cys355, Cys356, and Cys358) in the fourth intracellular loop (ICL4) are replaced with glutamines, adjacent to the hydrophobic LI350,352 motif on ICL4 (Figure 3B), which was previously confirmed^10^ to confer partial resistance to Nef_97ZA012_. Notably, this cysteine cluster is present in SERINC5 and SERINC4 and absent in other SERINC paralogs (Figure 3B). These findings further support the notion that the extended intracellular loop of SERINC5 plays a critical role in determining sensitivity to Nef-mediated downregulation.

### SERINC5 susceptibility to downregulation by HIV-1 Nef requires the intact tri- cysteine motif within ICL4

We investigated the role of each cysteine residue, individually and in combination, in the downregulation of SERINC5 by Nef. As shown in Figures 4A and 4B, single substitutions of Cys355, Cys356, or Cys358 minimally impaired the sensitivity of SERINC5 to Nef- mediated downregulation, while two (CC356,358QQ) or three cysteines caused an increasingly stronger impairment of downregulation, suggesting that the presence of all three cysteines simultaneously is required for maximum susceptibility to Nef. Given their proximity to the hydrophobic LI350,352 motif, we also examined whether simultaneous disruption of these two motifs would produce a synergistic effect. SERINC5 variants with either the LI350,352AA or the CCC355,356,358QQQ substitutions were downregulated by Nef to approximately 17% and 45% of baseline levels, respectively. However, LI350,352AA did not further reduce the downregulation of SERINC5 when combined with CCC355,356,358QQQ, indicating that the effects of these mutations are not additive.

**Figure 4:**
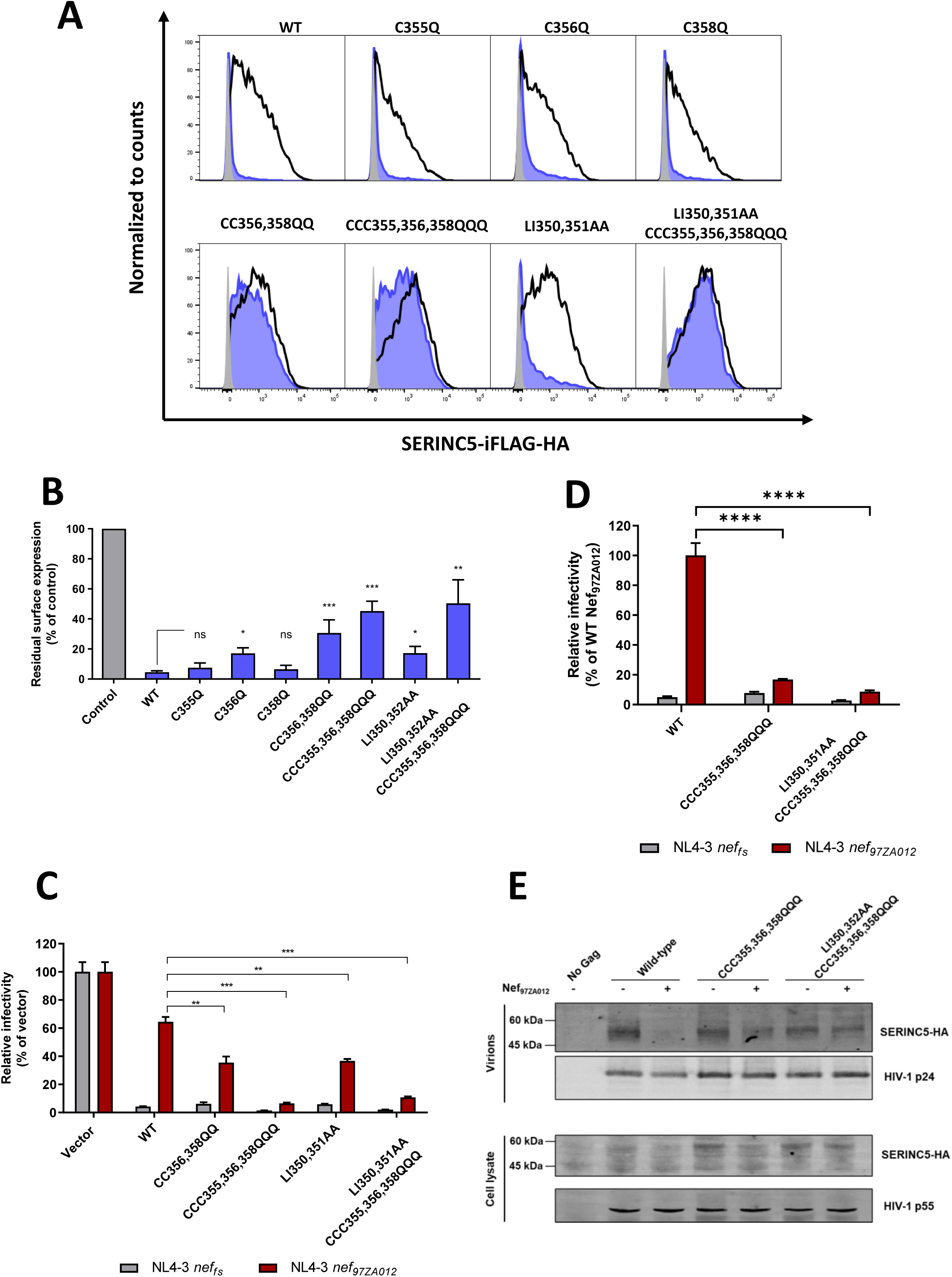
An intact cysteine cluster within SERINC5 ICL4 is required for antagonization by Nef_97ZA012_. (A-B) Effect of the indicated SERINC5 mutations on internalization efficiency by Nef_97ZA012_.S3/S5 DKO HEK293T cells were co-transfected with a SERINC5-iFLAG-HA variant, expressed from pcDNA3.1, and pIRES2-eGFP-Nef_97ZA012_ or empty vector control. SERINC5 surface expression in transfected (GFP^+^) cells was measured as median fluorescence intensity. (A) Representative flow cytometry profiles of SERINC5 variants in the presence of empty vector control (black line) or Nef_97ZA012_ (red area). The background signal is indicated in grey. (B) Residual surface expression of SERINC5 variants in the presence of Nef_97ZA012_, expressed as a percentage of the empty control sample. Bar graphs represent the mean + SD of n=3 biological independent experiments. Statistical significance was measured with an unpaired two-tailed t-test. *p<0.01; **p<0.001; ***p<0.0001; ns: non-significant; (C-E) Effect of the indicated SERINC5 mutations on HIV-1 infectivity and antagonization by Nef_97ZA012_. (C) Δenv HIV-1_NL4-3_ virions, harbouring either nef_97ZA012_ or nef_fs_, were produced in S3/S5 DKO HEK293T cells, co-transfected with PBJ5-Env_HXB2_ and a PBJ6- SERINC5-iFLAG-HA variant or empty vector control. Virus infectivity was measured on TZM-ZsGreen cells and is expressed as a percentage of the infectivity produced in the absence of SERINC5. A representative experiment of n=3 biological independent replicates is shown. Bar graphs indicate the mean + SD of n=4 technical replicates. Statistical significance was measured with unpaired two-tailed t-test. **p<0.001; ***p<0.0001; (D-E) Nef_97ZA012_ or nef_fs_ Δenv HIV-1_NL4-3_ were produced in S3/S5 DKO JTAg cells by electroporation with PBJ5-Env_HXB2_ and the indicated BPJ5-SERINC5-iFLAG-HA variant. (D) RT-normalized virus infectivity measured on TZM-ZsGreen cells, expressed relative to the infectivity of Δenv nef_97ZA012_ HIV-1_NL4-3_ in the presence of wild-type SERINC5. Bar graphs represent the mean + SD of n=4 technical replicates. Statistical significance was measured with unpaired two-tailed t-test. **p<0.001; ***p<0.0001; (E) Western Blot of SERINC5 variants in purified virus pellets and virus-producer cell lysates.

Next, we tested whether the role of Cys355, Cys356, and Cys358 in modulating SERINC5 downregulation by Nef correlates with the ability to rescue HIV-1 infectivity. As shown in Figure 4C, all tested variants inhibited HIV-1 effectively (10–20-fold), similar to the wild- type protein. While Nef_97ZA012_ counteracted wild-type SERINC5 by enhancing virus infectivity 16-fold, the rescue of infectivity by Nef was maximally reduced for the CCC355,356,358QQQ variant, consistent with the extent of downregulation observed in flow cytometry assays (Figure 4A and 4B).

Finally, we assessed whether the diminished ability of Nef_97ZA012_ to rescue HIV-1 infectivity in the presence of SERINC5 variants correlates with their levels of incorporation into virions. To this end, viral particles were produced by ectopically expressing SERINC5 variants in S3/S5 DKO Jurkat Tag cells instead of HEK293T cells, as the latter were previously found to yield virions contaminated with non-viral vesicles^2^, complicating the assessment of virally associated SERINC5. As observed with virus produced in HEK293T cells, Nef_97ZA012_ counteracted the inhibitory activity of wild-type SERINC5 in Jurkat Tag cells, enhancing infectivity ∼25-fold. However, Nef failed to overcome the inhibition imposed by the CCC355,356,358QQQ and LI350,352AA/CCC355,356,358QQQ variants (Figure 4D). This finding mirrors the incorporation data, where wild-type SERINC5 is efficiently excluded from virions in the presence of Nef_97ZA012_ (Figure 4E), while the CCC355,356,358QQQ and the LI350,352AA/CCC355,356,358QQQ variants are not.

These results confirm the critical role of the cysteine residues within the ICL4 region for the susceptibility of SERINC5 to Nef.

### The tri-cysteine motif is required for SERINC5 downregulation by lentiviral *nef* alleles, but is dispensable for Mo-MLV glycoGag and EIAV S2

Next, we explored whether the tri-cysteine and the hydrophobic LI350,352 motifs are required for SERINC5 downregulation by divergent *nef* alleles from HIV-1_LAI_ and HIV-1_SF2_ (both subtype B), SIV_Mac239_, SIV_Agm_, and by MoMLV glycoGag and EIAV S2. As in previous experiments, SERINC5 surface expression in S3/S5 DKO HEK293T cells was measured by flow cytometry following transfection. All retroviral antagonists efficiently internalized wild-type SERINC5, reducing its plasma membrane localization by more than 80% (Figure 5A-G). Among these, Nef_LAI_ was the least effective, leaving 20% residual surface expression (Figure 5B).

**Figure 5:**
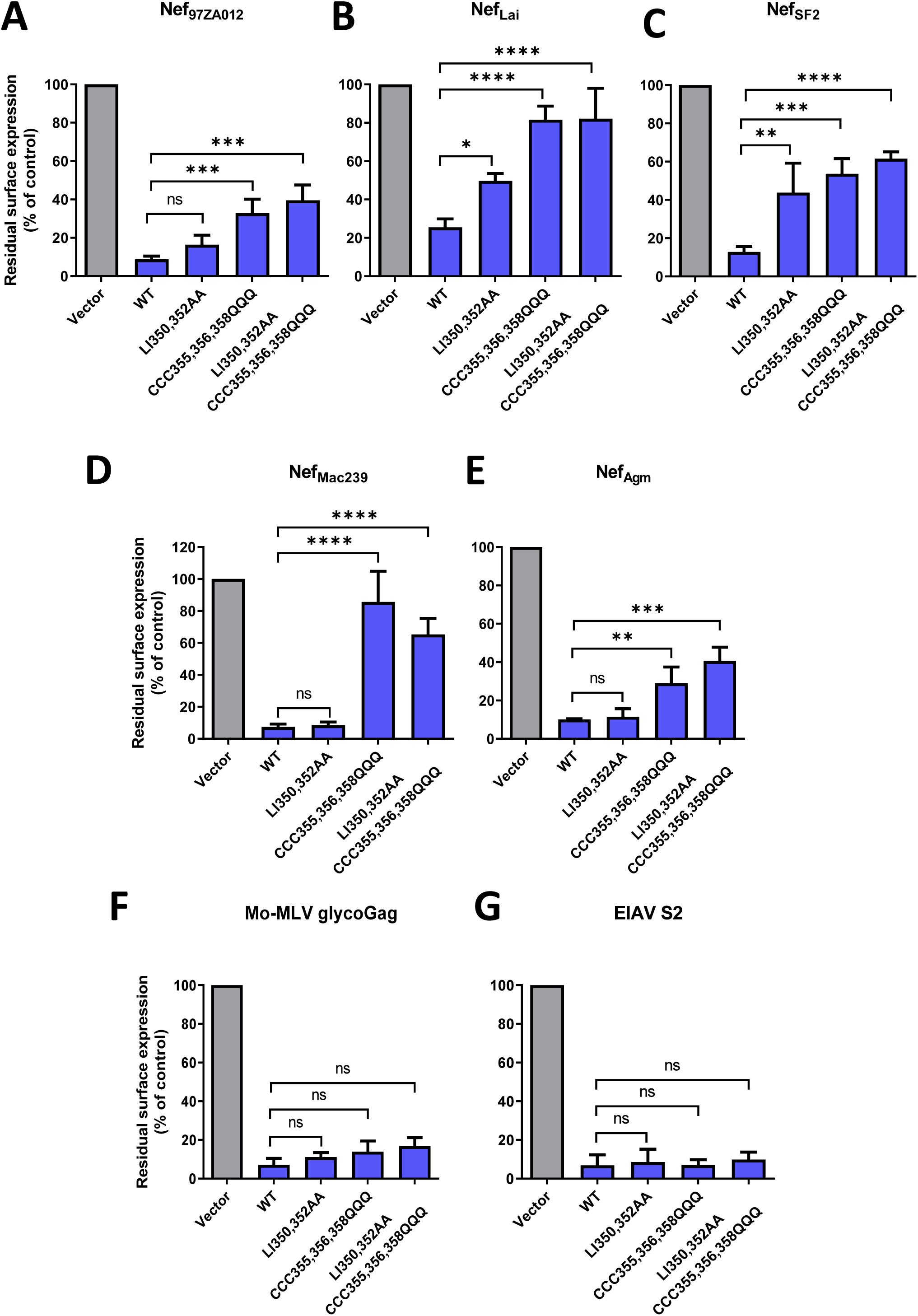
The Cysteine stretch within SERINC5 ICL4 is essential for internalization by lentiviral Nef, but dispensable for downregulation by MoMLV glycoGag and EIAV S2. Residual surface expression measured by flow cytometry of SERINC5 variants in the presence of diverse HIV-1 (A-C) and SIV (D-E) Nefs, MoMLV GlycoGag (F) and EIAV S2 (G). (A-G) S3/S5 DKO HEK293T cells were co-transfected with a SERINC5-iFLAG-HA variant expressed from PBJ5 and pIRES2-eGFP encoding a retroviral factor. SERINC5 residual surface expression was calculated as the median fluorescence intensity (MFI) in the presence of a retroviral factor relative to the MFI measured in the empty control sample, considered as 100% (grey bar). Bar graphs represent the mean + SD of n=3 biologically independent experiments.

Disruption of the CCC355,356,358 motif impaired SERINC5 downregulation by all *nef* alleles tested, although the degree of impairment varied, with downregulation by Nef from HIV-1_LAI_ and SIV_Mac239_ being the most affected (almost complete loss of downregulation) and Nef from HIV-1_SF2_ and SIV_Agm_ being the least (40% residual surface expression). By contrast, substitution of LI350,352 had little or no effect on SERINC5 susceptibility to Nef_97ZA012_, Nef_Agm_, or Nef_Mac239_. However, the LI350,352AA mutation did impair SERINC5 downregulation by *nef* alleles derived from HIV-1 subtype B LAI and SF2. Notably, combining LI350,352AA with CCC355,356,358QQQ did not result in increased impairment of SERINC5 downregulation by Nef_97ZA012_. Remarkably, neither the tri-cysteine nor the hydrophobic LI350,352 motif was crucial for SERINC5 internalization by MoMLV glycoGag and EIAV S2, as both retroviral factors reduced the surface expression of wild-type and mutant SERINC5 proteins similarly, to less than 5% of baseline levels (Figure 5F-G).

Overall, our results indicate that the cysteine stretch within ICL4, rather than LI350,352, is a common requirement for SERINC5 downregulation by all HIV-1 and SIV *nef* alleles tested. By contrast, SERINC5 downregulation by MoMLV glycoGag and EIAV S2 does not depend on either LI350,352 or CCC355,356,358, suggesting that the different retroviral factors have evolved to exploit distinct molecular features to counteract SERINC5.

### Cysteines 355,356,358 of SERINC5 are palmitoylated

Cysteine residues are targets of post-translational modifications, in particular S-palmitoylation, frequently detected in membrane proteins (reviewed in^25^). To explore whether SERINC5 is subject to this modification, we first examined potential palmitoylation sites *in silico* using the GPS-Palm algorithm. The analysis predicted multiple high-confidence cysteine targets in both SERINC3 and SERINC5, including several within the N-terminal cytoplasmic region and, for SERINC5, the tri-cysteine cluster (C355, C356, C358) located in intracellular loop 4 (ICL4).

To experimentally assess palmitoylation occupancies of SERINC3/5 cysteine residues, we employed a two-step differential labelling strategy using chemically equivalent isotopic variants of N-ethylmaleimide (NEM), the light (H_5_) and heavy (D_5_) forms, as thiol alkylating reagents, followed by mass-spectrometric quantification. Jurkat cell lines stably overexpressing twin-Strep-tagged versions of human SERINC3 and -5 were first established, and SERINC proteins were isolated by affinity chromatography. Proteins were first treated with H_5_-NEM to block free (unmodified) cysteine residues. Subsequently, thioacylated cysteines were hydrolysed by hydroxylamine, removing attached palmitoyl groups and exposing the previously modified cysteine thiols. These newly liberated thiols were then labelled with D_5_-NEM. To maximise sequence coverage, the proteins were digested with two proteases (trypsin and chymotrypsin) prior to liquid chromatography-tandem mass spectrometry (LC-MS/MS). Finally, the D_5_-NEM/H_5_-NEM labelling ratios of the resulting peptides were calculated to determine the extent of site-specific palmitoylation. Palmitoylation events were detected within the N-terminal peptides of SERINC3 (Figure 6A and extended data). By contrast, the N-terminal peptides of SERINC5 were not detected; therefore, palmitoylation in this region could not be assessed, preventing us from evaluating the predictions. However, robust palmitoylation (high stoichiometry and confidence) was confined to residues 355CCFCF359 within intracellular loop 4 (ICL4) of SERINC5 (Figure 6A and extended data).

**Figure 6:**
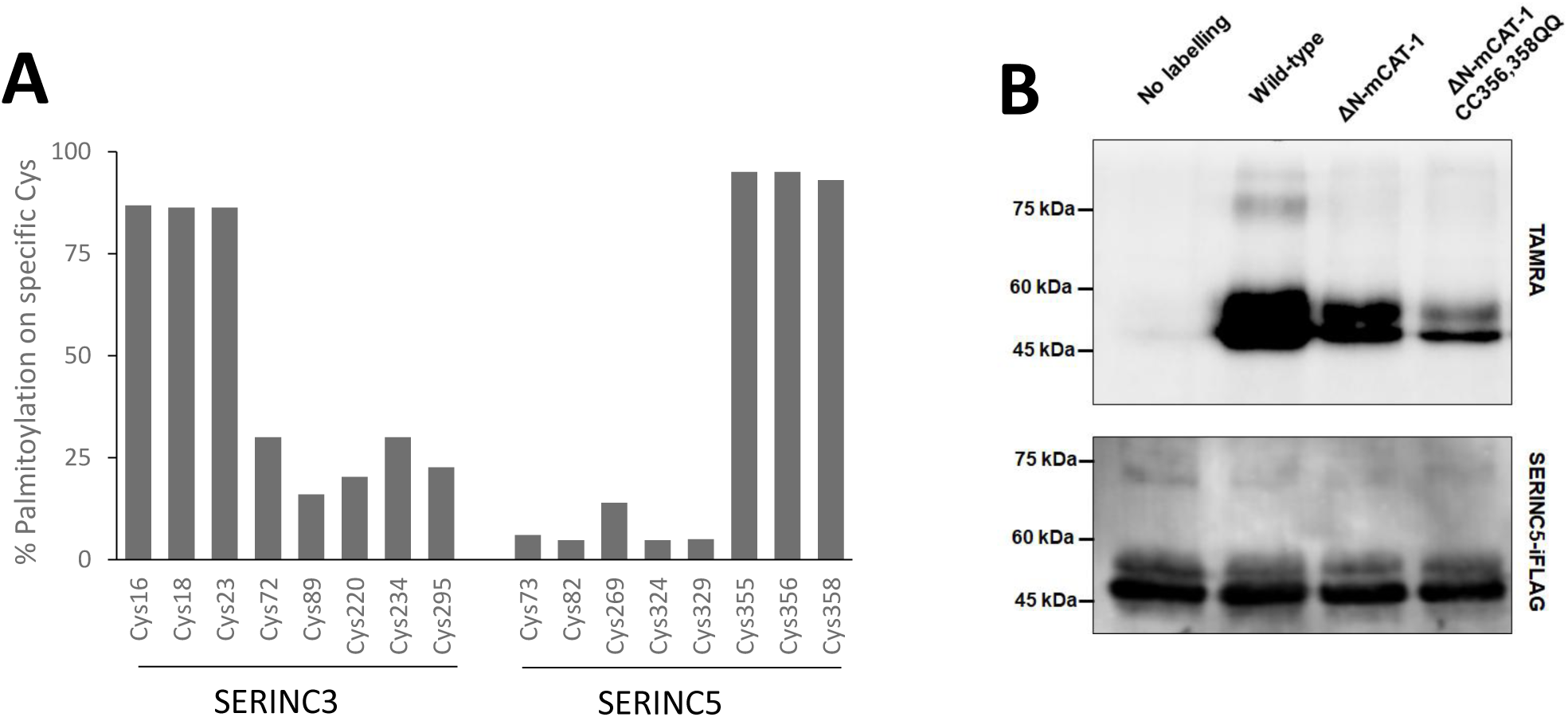
The cysteine stretch within SERINC5 ICL4 is palmitoylated. A) Percentage of palmytoylation of specific cysteines as computed from mass spectrometry acquisition of the D5-NEM/H5-NEM ratio of tryptic/chymotryptic peptides originated from SERINC5 (see extended data) (B) Detection of palmitoylation in SERINC5 by Click-chemistry. HEK293T cells were transfected with the indicated SERINC5-iFLAG variants and metabolically labelled by a clickable palmitate analogue, which was further modified by a TAMRA-labelled azide moiety. TAMRA and iFLAG were detected in cell lysates by Immunoblot.

To verify the SERINC5 results obtained by mass spectrometry, we employed a click-chemistry-based metabolic labelling approach. Cells expressing SERINC molecules were metabolically labelled with an alkynyl palmitate to enable copper-catalysed azide–alkyne cycloaddition chemistry. Because our mass spectrometry-based methodology could not assess palmitoylation of N-terminal cysteines, despite strong prediction by GPS-Palm, we specifically examined this region by engineering a SERINC5 variant in which the native N-terminal tail was replaced by the tail derived from the mCAT-1 transporter (ΔN- mCAT-1). To specifically assess the palmitoylation of CC356,358QQ, the ΔN-mCAT-1 variant carrying the cysteine to glutamine substitutions in positions 356 and 358 (ΔN- mCAT-1/CC356,358QQ) was also generated. All SERINC5 variants, containing also the internal FLAG, were individually overexpressed in HEK293T cells. Transfected cells were metabolically labelled with an alkynyl palmitate and SERINC5 molecules purified from cell lysates. Following SDS–PAGE, SERINC5 was detected via its FLAG epitope, and palmitoylation was visualized by in-gel fluorescence after ligation to a TAMRA-derivatized azido capture reagent. The experiment confirmed robust incorporation of alkynyl palmitate in wild-type SERINC5 (Figure 6B). Deletion of the N-terminus tail reduced the incorporation of metabolic labelling, suggesting that residues in this region are palmitoylated as predicted *in silico*. Further reduction of metabolic labelling was observed with the CC356,358QQ variant, confirming that these region of SERINC5 is also targeted by palmitoylation, in agreement with our LC–MS/MS data. Notably, TAMRA labelling was not completely prevented by the combined modifications, likely because of the presence of other minor palmitoylation sites, as predicted by GPS-Palm.

In conclusion, these results demonstrate that cysteines within ICL4, as well as residues at the N-terminus of SERINC5, are sites of palmitoylation. The partial discrepancy between the two detection methods likely reflects the limited recovery of N-terminal peptides during LC-MS/MS analysis, which can hinder identification of labile or hydrophobic modifications such as palmitoylation.

## Discussion

SERINC5 was discovered as a potent inhibitor of HIV-1 infectivity, effectively counteracted by multiple viral antagonists^2,3,17,18,26^. However, it exhibits several features that distinguish it from the canonical retroviral restriction factors, such as APOBEC3, TRIM5, and BST2. Indeed, SERINC5 does not appear to have evolved under positive selective pressure^27^, and its expression is not modulated by type I interferons^2^, both hallmark traits of many antiviral restriction factors. Moreover, its antiviral significance remains debated since SERINC5 fails to inhibit viral strains carrying certain envelope glycoproteins, including those from various primary HIV-1 isolates^28,29^ and other retroviruses^17,18^. These observations raise questions about the role of its antiviral activity.

Nonetheless, the evolutionary persistence of SERINC5 antagonists across three retroviral lineages^2,3,17,18^ and in SARS-CoV-2^19^ underscores its biological importance. This persistent evolutionary drive suggests that SERINC5 plays a role strong enough to impact viral evolution. In addition to SERINC5, other SERINC paralogs have shown varying degrees of antiviral activity. For example, SERINC3 was originally found to associate with virion particles and inhibit infectivity, though less potently than SERINC5^2,3^. These observations raise fundamental questions: what is the antiviral significance of the different SERINC paralogs? Have all these proteins influenced retroviral evolution? Do known viral antagonists target all SERINC paralogs equally, or is their activity specific to SERINC5?

Our results confirm that SERINC5 is the most potent inhibitor of HIV-1 infectivity among the SERINC family, with SERINC3 and SERINC1 showing moderate activity and SERINC2 being largely inactive. Interestingly, despite differences in antiviral potency, all SERINC paralogs are incorporated into HIV-1 virions. This suggests a shared mechanism of incorporation but differing impact on viral infectivity. Moreover, we confirmed that the ability of Nef to counteract SERINC proteins correlates with its ability to exclude them from virion particles and downregulate their surface expression in virus-producing cells.

Crucially, our data show that the breadth of the activity of different *nef* alleles and MoMLV glycoGag is selective for SERINC5. Except for EIAV S2, which downregulates SERINC3 as strongly as SERINC5, all viral factors have minimal or no activity against SERINC1, SERINC2 and SERINC3, but they efficiently target SERINC5. This reveals a fine-tuned counteraction that highlights an evolutionary adaptation to neutralize the most potent restriction factor, suggesting that SERINC5, rather than other paralogs, has been the main selective pressure driving the evolution of these viral antagonists.

The selective targeting of SERINC5 by viral factors indicates the presence of unique molecular features in SERINC5 that distinguish it from other paralogs. In this work, we identified a cluster of cysteine residues (Cys355, Cys356, and Cys358) within the fourth intracellular loop (ICL4) of SERINC5 as critical for its downregulation by all *nef* alleles tested. Interestingly, while the LI350,352 motif adjacent to the cysteine cluster also plays a role in downregulation, its effect is allele-specific, highlighting a complex interaction and different requirements within various viral isolates.

Although bimolecular fluorescence complementation assays have suggested a direct interaction between Nef and SERINC5^16,24,30^, definitive biochemical evidence supporting this notion is still missing. We hypothesize that the cysteine cluster may contribute to an interaction surface with Nef or alternatively, regulate the conformational state of ICL4 required for recognition. Mass spectrometry analysis demonstrated palmitoylation of these cysteines, suggesting that they may facilitate recognition by Nef indirectly, by maintaining the correct local conformation, rather than serving as direct contact residues. The observation that glycoGag and S2 retain full activity against SERINC5 cysteine mutants further supports the notion that different retroviral antagonists exploit distinct structural features of SERINC5 to achieve the same functional outcome.

In conclusion, this study provides evidence that SERINC5 is the primary target of viral antagonists among human SERINC paralogs and establishes the palmitoylated tri- cysteine motif within ICL4 as a critical determinant for its antagonism by Nef. Several questions remain to be addressed. Given that viruses can bypass SERINC5 inhibition by evolving resistant envelope glycoproteins, what forces retroviruses to retain a dedicated SERINC5-counteracting function? What is the structural basis of SERINC5 engagement by different viral proteins, and do these mechanisms converge at the level of endocytic machinery recruitment? Finally, what are the broader physiological roles of SERINC proteins in cellular processes beyond viral restriction?

Future research should explore these questions, with particular attention to the structural and functional dynamics of SERINC5 in the context of viral antagonism. Such studies may not only clarify the role of SERINC5 in the evolutionary host-virus arms race but could also identify opportunities to develop antiviral strategies that stabilize SERINC5 at the plasma membrane or interfere with its recognition by viral proteins.

## Materials and Methods

### Plasmids and reagents

The HIV-1 proviral plasmid *env*-deficient pNL4-3^2^, was modified with the insertion of NotI and XbaI unique sites flanking *nef* orf to allow the swap of Nef_NL4-3_ with desired variants. Specifically, here, *nef_NL4-3_* sequence was replaced with *nef_97ZA012_*. The HIV-1 proviral constructs were complemented with HIV-1_HXB2_ Env expressed from PBJ5 plasmid.

C-terminally HA-tagged SERINCs, used for infectivity and western blotting, have been previously generated^2^ and subcloned in pcDNA3.1, PBJ5 and PBJ6 plasmids. PCDNA3.1-mCAT-1 iFLAG encodes the murine cationic amino acid transporter 1 (mCAT- 1) modified by the insertion of a FLAG-tag within the third extracellular loop for the detection of surface expression by flow cytometry.

For flow cytometry analysis, Nef, glycoGag and S2 were cloned in pIRES2-eGFP, while the sequences of human SERINC-HA paralogs were further modified by PCR to incorporate a FLAG tag within the fourth extracellular loop (SERINC-iFLAG-HA). SERINC5 mutants were previously generated by site-directed mutagenesis^8^.

Mouse anti-HA (clone 16B12) was purchased from Biolegend, while mouse anti-FLAG (clone M2) was obtained from Sigma-Aldrich. The mouse anti-HIV-1 p24/p55 (Arp324) was obtained from the Centre of AIDS Research (CFAR). The secondary antibody goat anti-mouse IgG-IRDye680 was purchased from Li-COR. The secondary antibody goat anti-mouse IgG conjugated with APC was purchased from Jackson Immunoresearch.

### Cell lines

HeLa-TZM-zsGreen cells are derivative of the HeLa-TZM-bl HIV indicator cells (NIH AIDS Research and Reference Reagent program), modified to carry ZsGreen under the control of the HIV-1 LTR promoter, as previously described^1^. A SERINC3/5 double knock-out (S3/S5 DKO) HEK293T cell line was generated by transient transfection of PX458 and pLentiCRISPR v2 vectors, encoding Cas9 and a guide RNA targeting SERINC5 exon 2 (5’-CACCGGCTGAGGGACTGCCGAATCC-3’).

S3/S5 DKO HEK293T cells and HeLa-TZM-zsGreen cells were cultured in DMEM supplemented with 10% FBS and 2 mM L-Glutamine.

S3/S5 double knock-out Jurkat TAg (S3/S5 DKO JTAg) cells have been previously described^2^ and were cultured in RPMI complemented with 10% FBS and 2 mM L-Glutamine.

All cell lines were maintained in a humidified incubator at 37°C and 5% CO2.

### Infectivity assay

The effect of human SERINC paralogs on HIV-1 infectivity was investigated using virions limited to single-cycle replication by transfecting S3/S5 DKO HEK293T cells. For each sample, 3 million cells were seeded in a 10 cm plate, the day before transfection. 15 µg of an *env*-defective HIV-1_NL4-3_ proviral plasmid, encoding an inactive *nef* or the clade C *nef_97ZA012_* allele, were complemented with 2 µg of PBJ5-Env_HXB2_ and co-transfected with either 5 µg of PBJ6 or 1 µg of PBJ5, expressing a SERINC paralog. Transfection was performed by calcium-phosphate co-precipitation. Forty-eight hours post-transfection, virus-containing supernatants were harvested, clarified by centrifugation (300g for 5 min) and filtered with 0.22 µm filters. The amount of virus produced in cell supernatants was quantified by SG-PERT reverse transcription assay, as previously described^31^. Virus infectivity was assessed by inoculating five serial dilutions onto HeLa-TZM-ZsGreen cells in 96-well plates. Infection was performed in quadruplicate and scored with the Ensight plate reader (Perkin Elmer). The infectivity of each virus was calculated by normalising the number of infected cells to the respective RT-activity. For each sample, the infectivity was then expressed relative to the infectivity of the virus in the absence of SERINC, which was considered to be 100%.

### Analysis of virion-incorporated human SERINC paralogs

To detect virion-associated SERINC proteins, HIV-1_NL4-3_, encoding *nef_fs_* or *nef_97ZA012_* and limited to single cycle replication, was produced by transfection of S3/S5 DKO HEK293T, as described above (infectivity assay). Forty-eight hours after transfection, virus-producing cells were collected in PBS and lysed for 30 min in an ice-cold buffer containing 100 mM NaCl, 10 mM HEPES pH 7.5, 1 mM TCEP pH 7 (Sigma), 1% n-Dodecyl beta-D-maltoside (Sigma) and 2X protease inhibitor cocktail (Roche). Lysates were clarified by centrifugation (16,000g for 20 min at 4°C) and supplemented with Laemmli Buffer, containing 50 mM TCEP. In parallel, virus-containing cell supernatants were clarified by centrifugation, filtered through a 0.22 µm membrane, overlaid on a 25% sucrose cushion and purified by ultracentrifugation at 100,000g for 2 hours (SW41 rotor, Beckman Coulter). The concentrated virus pellets were resuspended in Laemmli Buffer, supplemented with 50 mM TCEP. Samples were separated on 12.5% acrylamide tricine gels and transferred onto an Immobilon-FL PVDF membrane (Millipore). Membranes were probed with mouse anti-HA antibody (HA.11 clone 16B12, diluted 1:500) and mouse anti-HIV-1 p24/p55 antibody (Arp324, diluted 1:500). For detection, a goat anti-mouse IgG IRDye 680 was used, diluted 1:5000. Revert Total protein staining (LI-COR) was used post-acquisition to normalize the amount of sample loaded. Blots were imaged using Odyssey infrared imager (LI-COR).

### Analysis of SERINCs surface downregulation by retroviral antagonists

The sensitivity of human SERINC proteins to cell surface downregulation by retroviral antagonists was monitored by flow cytometry. S3/S5 DKO HEK293T cells were seeded in a 24-well plate at a density of 150.000 cells/well. Transfection was performed by calcium-phosphate co-precipitation. Cells were co-transfected with 50 ng of pcDNA3.1- SERINC-iFLAG-HA or 100 ng of PBJ5-SERINC-IFLAG-HA, and 300 ng of pIRES2-eGFP plasmid expressing a retroviral factor or empty vector control. Two days post-transfection, cells were processed for flow cytometry staining: cells were collected, washed in PBA (PBS, 5% BSA, 0,1 % sodium azide) and incubated with anti-FLAG antibody (1:500 in PBA) 45 min at 4°C. Cells were then washed twice in PBA and stained with an APC-conjugated anti-mouse IgG (1:500 in PBA) for 45 min at 4°C. After washing twice with PBA, cells were fixed in 4% PFA (Sigma) and analysed with the 2-laser BD Canto flow-cytometer. Single viable cells were selected according to FSC-A vs SSC-A and then FSC-A vs FSC-H. Transfected cells were selected according to eGFP positivity. Acquisition gates were defined using FACSDiva software, while post-acquisition analyses were performed using FlowJo software. Residual surface expression was calculated as the amount of SERINC protein detected on the plasma membrane in the presence of a viral antagonist, relative to the surface levels measured in the presence of an empty vector control.

### Palmitoylation analysis

SERINC3 (NP_006802.1) and SERINC5 (NP_001167543.1) amino acid sequences were subjected to in silico prediction analysis of potential palmitoylation sites using GPS- Palm^32^.

The isolated SERINC proteins were precipitated with chloroform/methanol, washed with methanol, and the protein pellets were allowed to air-dry. Pellets were resuspended in 4% SDS, 1 mM EDTA, 150 mM NaCl, 50 mM triethanolamine (TEA), pH 7.4, and incubated with H5-NEM (25 mM final concentration) for 2 h at room temperature (RT). Proteins were again precipitated with chloroform/methanol, washed three times with methanol, air-dried, and resuspended in 4% SDS, 1 mM EDTA, 150 mM NaCl, 50 mM TEA, pH 7.4. Thioesters were hydrolyzed by treatment with hydroxylamine (NH₂OH, 0.75 M final concentration) for 2 h at RT. Proteins were precipitated with chloroform/methanol, washed three times with methanol, and air-dried. Pellets were resuspended in 4% SDS, 1 mM EDTA, 150 mM NaCl, 50 mM TEA, pH 7.4, and incubated with D5-NEM (25 mM final concentration) for 2 h at RT. Proteins were then precipitated with chloroform/methanol, washed three times with methanol, air-dried, and resuspended in HEPES buffer, pH 8.0, for digestion with sequencing-grade trypsin or chymotrypsin (Promega) for 2 h and 16 h. Digests corresponding to the same protease were combined, and the resulting peptides were desalted on C18 StageTips and dried using a SpeedVac concentrator. Prior to LC-MS/MS measurements, peptide samples were resuspended in 0.1% TFA and 2% acetonitrile in water. Chromatographic separation was performed on an Easy-Spray Acclaim PepMap column (50 cm × 75 µm i.d., Thermo Fisher Scientific) using a 70 min acetonitrile gradient in 0.1% aqueous formic acid at a flow rate of 250 nl/min. An Easy-nLC 1000 system was coupled to a Q Exactive mass spectrometer via an Easy-Spray source (all Thermo Fisher Scientific). The Q Exactive was operated in data-dependent mode, with survey scans acquired at a resolution of 70,000 at m/z 200. Up to 10 of the most abundant isotope patterns with charges 2–7 from the survey scan were selected with an isolation window of 2.0 m/z and fragmented by higher-energy collision dissociation (HCD) with normalized collision energy of 25. The dynamic exclusion was set to 20 s. The maximum ion injection times for survey and MS/MS scans (acquired at a resolution of 17,500 at m/z 200) were 20 and 120 ms, respectively. The ion target value was set to 1 × 10⁶ for MS and 1 × 10⁵ for MS/MS, with the minimum AGC target set to 1 × 10². Data were processed with PEAKS Studio 8.5 (Bioinformatics Solutions Inc.) and searched against the reference human proteome (UniProt). Specific cleavage by trypsin or chymotrypsin was selected, allowing up to two missed cleavages for trypsin and up to five for chymotrypsin. The maximum precursor mass error was set to 5 ppm and the fragment mass error to 0.01 Da. Methionine oxidation was set as a variable modification, and cysteine residues were set to be modified with either D_5_-NEM or H_5_-NEM. The maximum number of modifications per peptide was set to five. The peptide-level false discovery rate (FDR) was set to 0.01. The D_5_-NEM/H_5_-NEM ratios were calculated by the software; when only H_5_-NEM signal with no corresponding D_5_- NEM signal, or vice versa, was detected the ratio was set to 0.5 or 20, respectively. Final site occupancies were calculated as the arithmetic mean of D_5_-NEM/H_5_-NEM ratios across all peptides containing a given site.

Transfected cells were metabolically labelled with a “clickable” palmitate analogue - 17- Octadecynoic Acid; the cells were lysed, and the metabolically labeled proteins were ligated to an azido capture reagent derivatized with biotin and TAMRA, following a previously reported protocol^33^. SERINC5 proteins were purified from cell lysates and resolved by SDS–PAGE. Palmitoylation was detected by in-gel fluorescence, and SERINC5 variants were detected via the FLAG epitope by Western blotting.

### Statistical analysis

Statistical analysis was performed using GraphPad Prism. For infectivity assays, statistical significance was computed by performing an unpaired two-tailed t-test with Welch’s correction. Instead, the statistical analysis of SERINCs downregulation by retroviral antagonist was performed with one-sample t-test, considering 100 as hypothetical value. *p<0.05; **p<0.01; ***p<0.001; not significant.

## ACKNOWLEDGMENTS

This work was supported by MUR PNRR BaC PE INF-ACT S1 GENESIS and by PRIN 2022RYNKB5 to MP. The laboratory of PC is supported by the Francis Crick Institute, which receives its core funding from Cancer Research UK (CC2058), the UK Medical Research Council (CC2058), and the Wellcome Trust (CC2058). Authors made the following contributions: conceived and designed the experiments, MP, PC, RS, ET, CB, AC. performed the experiments, CB, AR, TV, AC, MB, RS, PC, MP. analyzed the data, CB, RS, ET, PC, MP; wrote the paper, CB, RS, ET, PC, MP.

